# Conserved metabolic vulnerabilities across pathogenic coronaviruses nominate host-directed therapeutic targets

**DOI:** 10.64898/2026.04.17.716662

**Authors:** Bushra Dohai, Diana C. El Assal, Mina Kang, Ashish K. Jaiswal, Christophe Poulet, Sarah Daakour, David R. Nelson, Pascal Falter-Braun, Jean-Claude Twizere, Kourosh Salehi-Ashtiani

## Abstract

Pathogenic coronaviruses profoundly rewire host cell metabolism to support viral replication, yet whether these metabolic alterations expose shared and actionable vulnerabilities remains unclear. By integrating transcriptomic profiles from cells infected with SARS-CoV, SARS-CoV-2, and MERS-CoV with genome-scale metabolic models, we identify conserved and virus-specific metabolic perturbations affecting mitochondrial transport, nucleotide biosynthesis, fatty acid metabolism, and redox balance. Despite distinct transcriptional responses, all three viruses converge on a limited set of metabolic reactions whose flux ranges deviate strongly from healthy states. Using a network-based predictive framework, we systematically identify gene-pair perturbations that restore perturbed reaction fluxes toward non-infected metabolic states. Predicted rescue mechanisms reveal shared metabolic dependencies across coronaviruses, as well as time-dependent virus-specific vulnerabilities, and nominate druggable host targets. Notably, several top predictions align with independent experimental and clinical evidence, including metabolic interventions shown to reduce viral replication or disease severity in COVID-19 patients. Together, our results define conserved metabolic rescue pathways in coronavirus infection and provide a general strategy for identifying host-directed therapeutic opportunities from transcriptomic data.

**Highlights:**

- Coronaviruses converge on shared metabolic vulnerabilities in host cells
- NiTRO predicts gene pairs that rescue viral-induced metabolic states
- Mitochondrial transport emerges as a key pan-coronavirus target
- Top predictions validated by clinical trials and *in vitro* evidence

**eTOC Blurb:** Dohai et al. develop NiTRO, a network-based algorithm that integrates coronavirus-induced transcriptomic changes with genome-scale metabolic models to identify gene-pair perturbations capable of rescuing infected metabolic states. The approach reveals shared and virus-specific druggable metabolic vulnerabilities, with top predictions corroborated by clinical evidence.

## INTRODUCTION

Current known human coronaviruses (HCoV) can be classified into two categories based on their pathogenicity. Seasonal viruses HCoV-229E,^1^ HCoV-OC43,^2^ HCoV-NL63,^3^ and HCoV-HKU1^4^ induce generally mild upper-respiratory tract symptoms and are responsible for up to 30% of common colds in adults.^5^ However, in infants, elderly and immunocompromised individuals, they are able to cause fatal lower-respiratory tract infections.^6^ The second category comprises highly pathogenic severe acute respiratory syndrome virus (SARS-CoV),^7^ SARS-CoV-2^8,9^ and Middle East respiratory syndrome (MERS-CoV),^10^ which are able to rapidly spread from the upper-respiratory epithelial cells to infect the lower-respiratory tract to cause severe diseases. Viruses in this category have the ability to cause epidemics and global pandemics such as COVID-19.^8,11^ Coronaviruses have the potential to mutate or recombine and generate new variants that escape vaccines.^12^ A number of direct-acting antivirals have been approved for COVID-19 treatment. These include Remdesivir, a drug approved to treat patients suffering from severe COVID-19 in 2020.^13,14^ Remdesivir mimics RNA nucleotides and terminates viral RNA chain elongation by the RNA-dependent RNA polymerase (RdRp).^15^ However, this inhibition is not optimal, and the treatment was not widely effective.^16,17^ Ultimately, the WHO recommended against the use of Remdesivir in November 2020.^18^ Another RdRp inhibitor, Molnupiravir, has a broad-spectrum antiviral effect for several CoVs, including SARS-CoV, CoV-2 and MERS-CoV,^19^ and remains an affordable and effective inhibitor against COVID-19 to this day.^20,21^ Additional inhibitors, ensitrelvir and nirmatrelvir, targeting the main protease enzyme of SARS-CoV-2 (Mpro) exhibit broad-spectrum antiviral activities and are approved to be used in clinics.^22^ However, the occurrence of drug-resistant mutations in the targeted viral enzymes has been the major obstacle against any direct-acting antiviral,^23–25^ and combinatorial therapies are already showing elevated efficacy in animal models for SARS-CoV-2.^26,27^ Alternative treatment strategies include targeting dependency host factors identified in CRISPR screens,^28–31^ host proteins that interact with CoV factors,^32–37^ and other cellular mechanisms such as phosphorylation^34,38,39^ and metabolic pathways.^40,41^ The latter is an important therapeutic target as it became clear already early in the COVID-19 pandemic that metabolic disorders including diabetes, hypertension, cardiovascular diseases and obesity constitute risk factors for severe symptoms and fatal COVID-19.^8,42–46^ At the molecular level, perturbations of cellular metabolism by SARS-CoV and SARS-CoV-2 coronaviruses are orchestrated, at least in part, through their direct binding to their receptor angiotensin converting enzyme-2 (ACE2), a negative regulator of the renin-angiotensin system.^47–50^ This binding down-regulates ACE2 from the cell membrane and impacts its basic function, which is primary degradation of angiotensin II to angiotensin1-7.^51^ As a result, excess angiotensin II molecules bind to angiotensin type 1 (AT1) receptors and induce detrimental effects such as vasoconstriction, fibrosis, inflammation, thrombosis and pulmonary damages.^51^ At the genome-wide level, metabolic modeling (GEM), using gene expression data on different SARS-CoV-2-infected cells or patient samples, has shown that it was possible to predict additional clinically relevant metabolic targets beyond ACE-2 by inferring and comparing metabolic fluxes between infected and non-infected cells.^40^ Thus, to identify metabolic drug targets for highly pathogenic betacoronaviruses, we have generated RNA-sequencing data from cells infected with SARS-CoV, SARS-CoV-2, and MERS-CoV, at different time courses. These data were then integrated with the human genome-scale metabolic model, Recon3D,^52^ to obtain six flux models of betacoronavirus-infected and non-infected cells. We designed a new tool, NiTRO, that extends metabolic transformation frameworks^53^ to combinatorial gene perturbations, and that (i) detects viral-induced metabolic changes, (ii) predicts the consequences on reaction fluxes following in silico double-gene deletions, and (iii) identifies novel metabolic drug-target pairs. We show that this strategy has the power to predict common and distinct host metabolic targets of the studied betacoronaviruses, and is broadly applicable to drug target identification against pathogens.

## RESULTS

### A comparative analysis of transcriptomic changes associated with SARS-CoV, SARS-CoV-2 and MERS infection

Highly pathogenic coronaviruses such as SARS-CoV, SARS-CoV-2, and MERS-CoV cause severe respiratory disease and share substantial genome sequence similarity (>50%).^54^ Infection by these viruses triggers large-scale host transcriptional reprogramming, affecting thousands of genes, as previously reported in multiple cellular and animal models.^55–59^

To characterize shared and virus-specific host responses, we infected HEK293-ACE2 cells with SARS-CoV, SARS-CoV-2, or MERS-CoV at a multiplicity of infection of 0.01 and performed RNA sequencing at 24 and 48 hours post-infection. At 24 hours, infection with SARS-CoV, SARS-CoV-2, and MERS-CoV resulted in significant differential expression of 2,823, 1,795, and 2,852 human genes, respectively (Figure 1A). Among these, 775 genes were commonly deregulated by all three viruses, while distinct subsets were uniquely affected by each virus.

**Figure 1.**
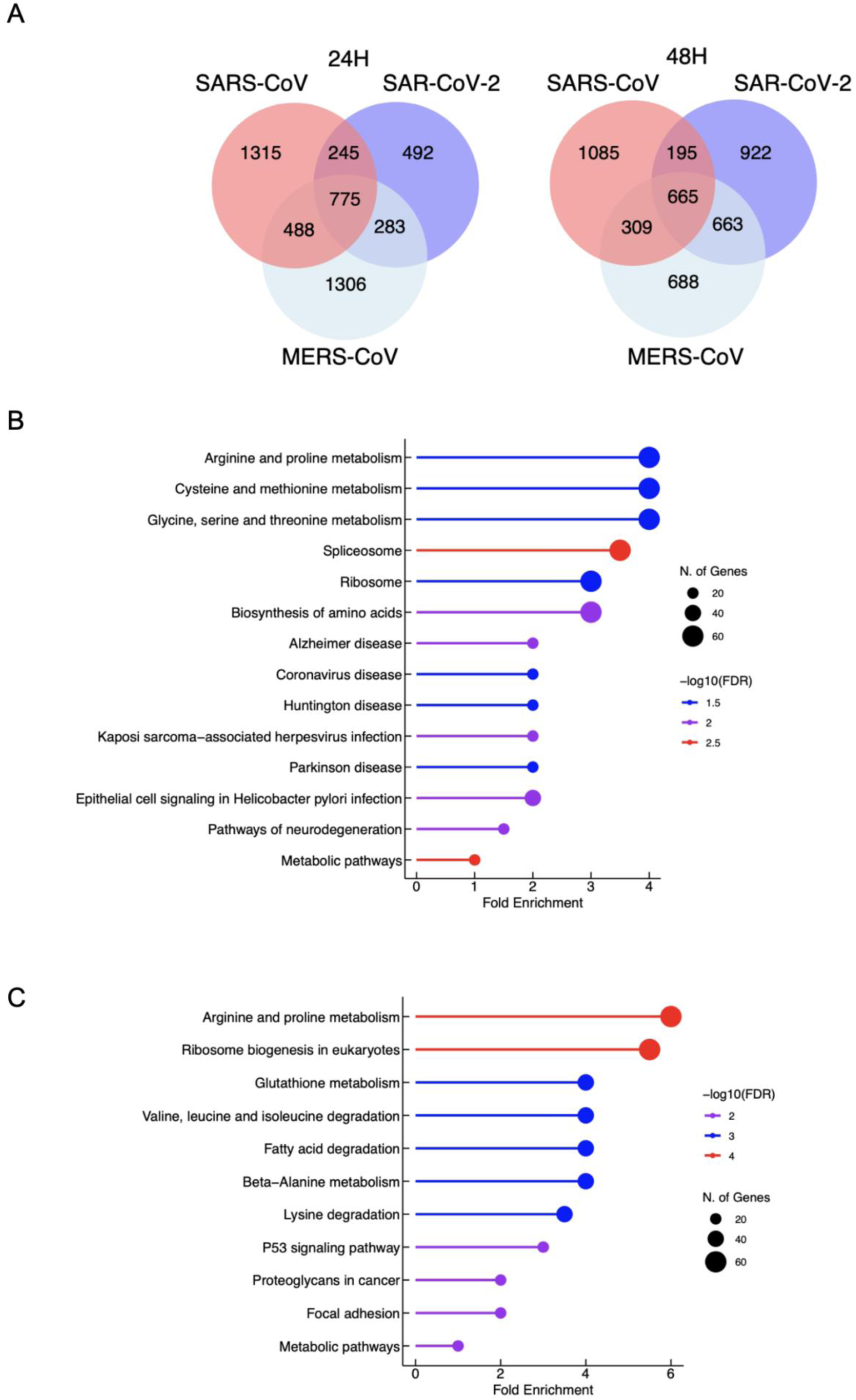
A comparative analysis of transcriptomic changes associated with SARS-CoV, SARS-CoV-2 and MERS infection. Venn diagrams showing the number of differentially expressed genes in HEK-293 cells at 24- and 48-hours post-infection by indicated coronaviruses. **(B)** and **(C)** Common metabolic processes enriched following infection with SARS-CoV, -CoV-2 and MERS at 24- and 48-hours post-infection, respectively.

At 48 hours post-infection, transcriptional perturbations remained extensive, with 2,254 genes altered in SARS-CoV–infected cells, 2,445 in SARS-CoV-2–infected cells, and 2,325 in MERS-CoV–infected cells (Figure 1A). Of these, 665 genes were commonly affected across all infections, while each virus also exhibited a large set of uniquely deregulated genes.

Gene set enrichment analysis revealed that, in addition to inflammatory and immune-related pathways (Figure S1) which has also been widely reported by others,^60–62^ genes involved in cellular metabolism were consistently enriched among the commonly deregulated genes at both time points (Figure 1B). These included pathways associated with amino acid metabolism, lipid metabolism, and nucleotide biosynthesis. These findings indicate that metabolic reprogramming is a conserved feature of infection by highly pathogenic coronaviruses.

### Coronaviruses increase biomass production and globally perturb metabolic fluxes

To assess how transcriptional changes translate into functional metabolic alterations, we integrated RNA-seq data from infected and non-infected cells with the human genome-scale metabolic model Recon3D^52^ using the GIMME algorithm.^63^ This approach yielded six virus-specific metabolic models corresponding to SARS-CoV, SARS-CoV-2, and MERS-CoV infection at 24 and 48 hours, along with two control models (uninfected, time points).

We then applied flux balance analysis (FBA)^64,65^ to compute reaction flux distributions and estimate biomass production as a proxy for cellular growth capacity. Comparison of infected and control models revealed virus-specific differences in biomass trends over time (Figure 2A, Supplementary Table 2A). SARS-CoV infection was associated with a modest decrease in biomass flux, SARS-CoV-2 infection resulted in increased biomass production, and MERS-CoV infection showed minimal change relative to controls (Figure 2A). Despite these differences, all infected models exhibited higher absolute reaction flux values compared to non-infected controls (Figures 2B–2E, Supplementary Table 2B). Consistent with this global increase in metabolic activity, infected models displayed a substantially larger number of perturbed reactions than controls (Figures 2F–2K). These results indicate that coronavirus infection broadly enhances metabolic throughput, likely to support viral replication demands, a phenomenon previously observed in other viral infections.^66,67^

**Figure 2.**
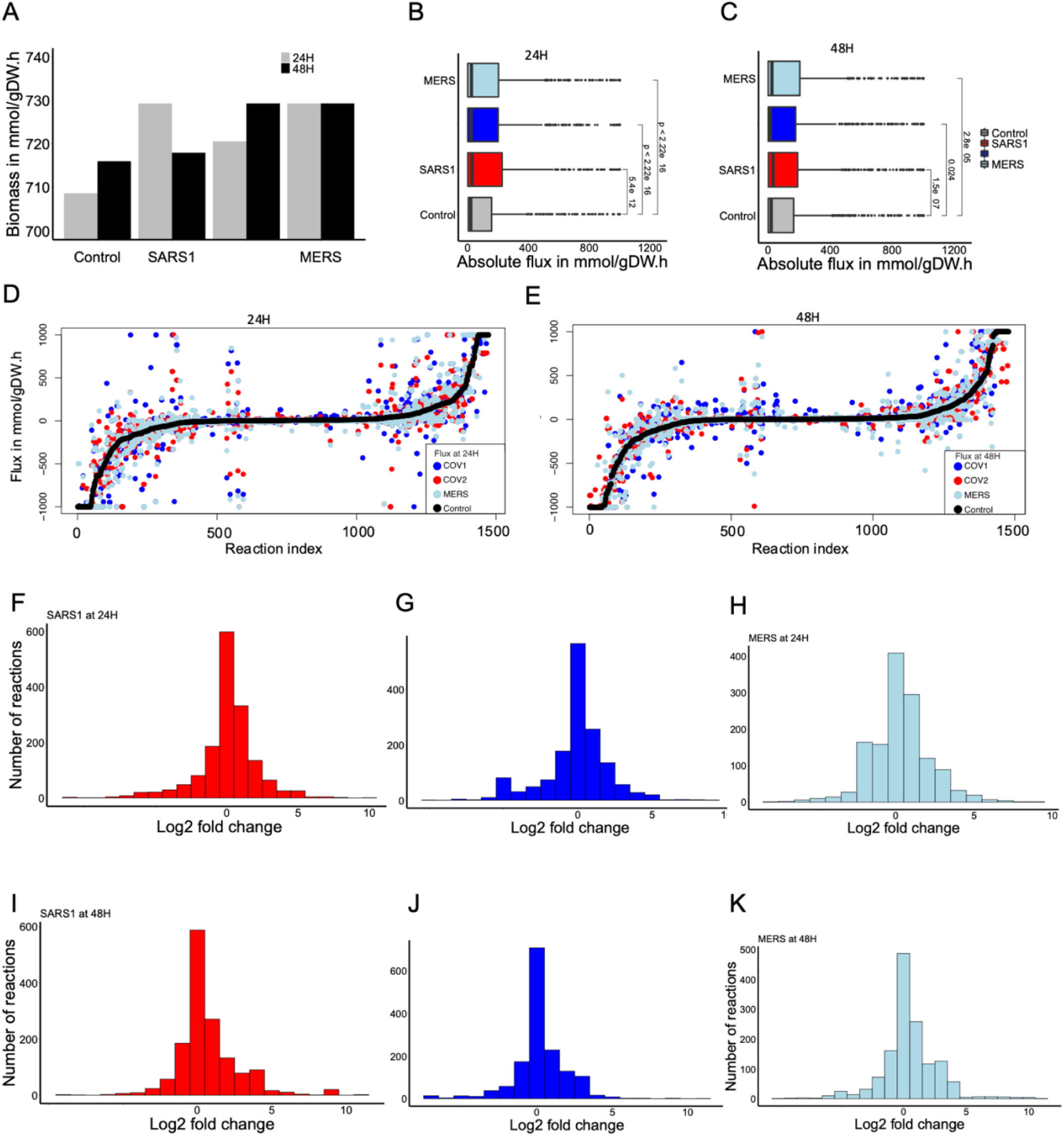
Coronaviruses increase cell biomass and induce metabolic perturbations. **(A)** The bar graph corresponds to the maximum biomass growth of the respective model (x-axis).

### Coronavirus infection perturbs key metabolic pathways

We next sought to identify metabolic pathways most strongly affected by coronavirus infection. A reaction was defined as perturbed if its flux in an infected model deviated by at least two-fold from the corresponding control model (Supplementary Tables 2C-2H). At 24 hours post-infection, 549, 545, and 510 reactions were perturbed in the SARS-CoV, SARS-CoV-2, and MERS-CoV models, respectively. At 48 hours, the number of perturbed reactions remained high, with 510, 414, and 492 reactions affected, respectively. Pathway enrichment analysis revealed that all three viruses consistently altered pathways related to mitochondrial and peroxisomal transport, fatty acid synthesis and oxidation, glycolysis, nucleotide metabolism, and branched-chain amino acid metabolism (Figure 3A, Supplementary Table 2I). In addition to these shared perturbations, virus-specific effects were observed. For example, oxidative phosphorylation and vitamin E metabolism were preferentially affected in SARS-CoV infection, while vitamin D metabolism was selectively perturbed in SARS-CoV-2 infection (Figure 3A). At 48 hours, cholesterol metabolism and fatty acid oxidation emerged as commonly perturbed pathways, while virus-specific alterations included oxidative phosphorylation (SARS-CoV), vitamin C metabolism (SARS-CoV-2), and sphingolipid metabolism (MERS-CoV) (Figure 3B). These results indicate that highly pathogenic coronaviruses induce both conserved and virus-specific metabolic rewiring, with mitochondria-centered pathways emerging as major targets.

**Figure 3.**
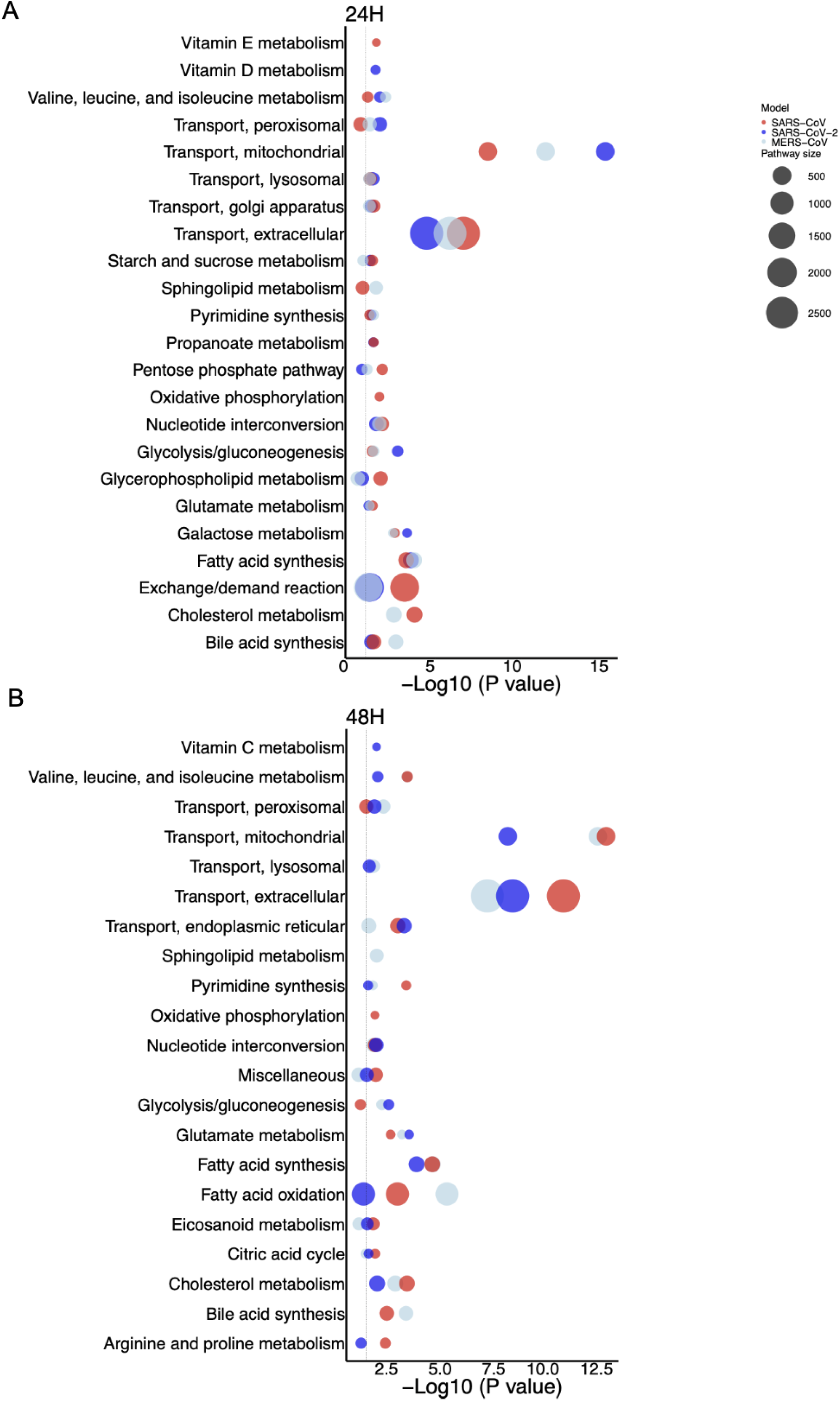
Functional enrichment analysis of the perturbed reactions. Pathways that are significantly (FDR < .05) enriched in virus-induced perturbed reaction sets.

### Infection alters metabolic reaction activation states

Beyond changes in flux magnitude, viral infection also altered the activation status of individual reactions. Reactions carrying non-zero flux were considered active, while those with zero flux were considered inactive. At 24 hours post-infection, 160 reactions exhibited altered activation status relative to control models (Supplementary Table 2K). Among these, all three viruses activated a shared set of reactions involved in nucleotide processing, folate metabolism, and amino sugar metabolism (Figure 4A). These activations are consistent with increased biosynthetic demand during viral replication. In contrast, 125 reactions were commonly deactivated across infections.

**Figure 4.**
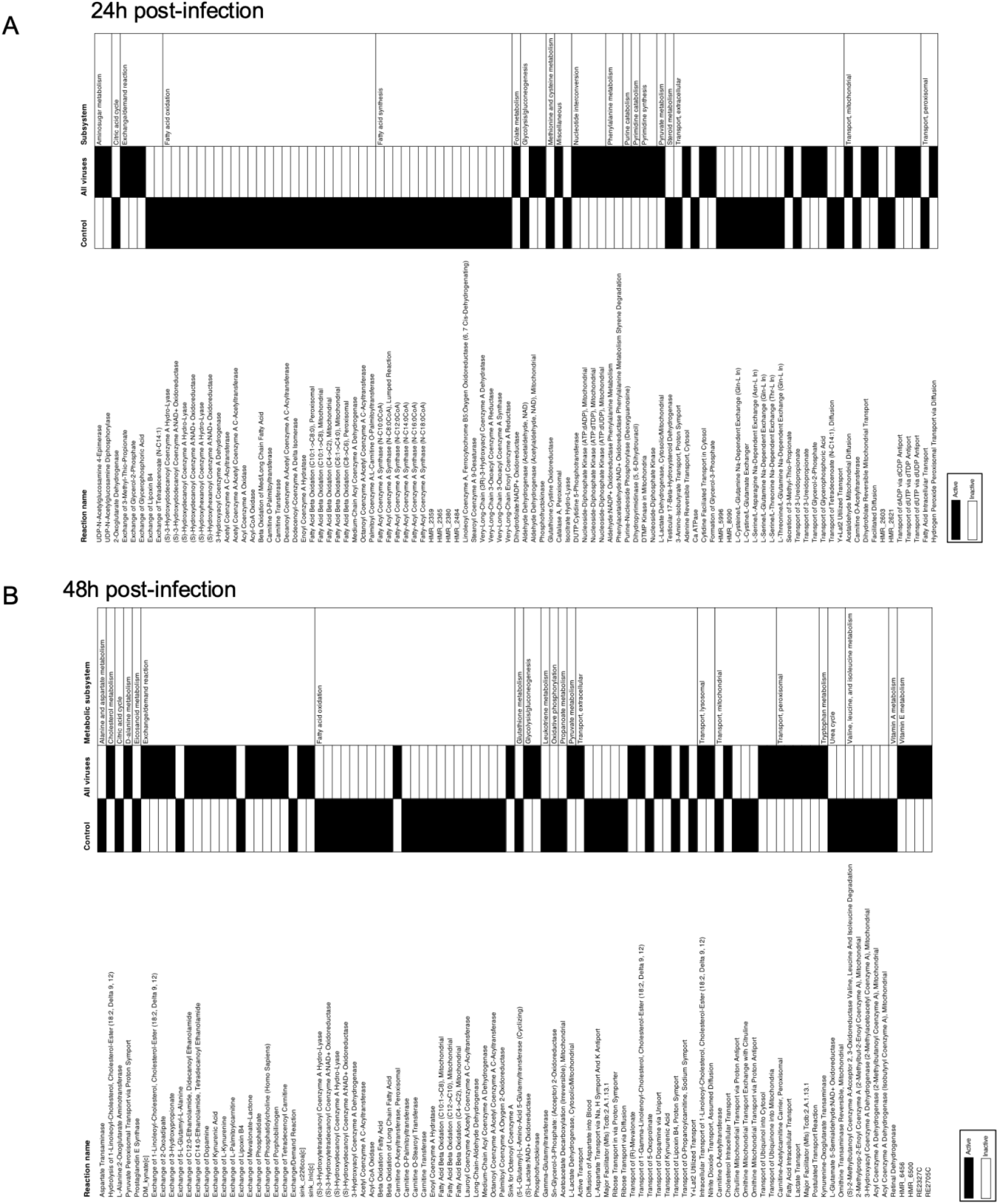
Perturbation on the reaction activation status. An active reaction is a reaction that carries non-zero flux (black box), and an inactive reaction is a reaction that carries zero flux (white box). In each panel, the first column lists the reactions in which the activation status was altered due to infection. The second column indicates the activation status of the corresponding reaction in the control model. The third column indicates the consensus activation status of the corresponding reaction in the infected models (SARS-CoV, -CoV2, and MERS-CoV). The fourth column lists the metabolic subsystem that is assigned to the corresponding reaction. (**A**) and (**B**) panels are for 24H and 48H models, respectively.

These reactions predominantly involved fatty acid metabolism, glycolysis, sulfur amino acid metabolism, and steroid metabolism (Figure 4A). Together, these shifts suggest a coordinated redistribution of metabolic resources rather than random reaction-level disruption. At 48 hours post-infection, 110 reactions displayed altered activation status (Supplementary Table 2L). A subset of reactions involved in cholesterol metabolism, tryptophan metabolism, and vitamin metabolism became activated, while reactions linked to amino acid catabolism, redox balance, and oxidative phosphorylation were deactivated (Figure 4B). These dynamic changes indicate that metabolic rewiring evolves over the course of infection.

### Directionality changes reveal altered metabolic equilibria

We next examined changes in reaction directionality, defined by the sign of reaction flux. A reversal in flux sign reflects a shift in reaction equilibrium and metabolic function. At 24 hours post-infection, many reactions involved in nucleotide metabolism exhibited consistent positive flux across infected models, indicating enhanced production of nucleotides (Figure S2A, Supplementary Table 2M). This observation aligns with the increased demand for nucleotides required for viral genome replication, as previously shown for other viruses.^68,69^ At 48 hours, nucleotide metabolism appeared more balanced, while reactions involving mitochondrial enzymes such as pyruvate kinase and pyruvate phosphotransferase showed increased activity (Figure S2B, Supplementary Table 2N). These findings are consistent with previous reports linking mitochondrial metabolism to disease severity and therapeutic response in viral infections.^70^

### Coronaviruses perturb mitochondria transport reactions

To rank metabolic reactions affected during coronavirus infection, we calculated fold changes in reaction fluxes for each infected model at 24 and 48 hours post-infection relative to the corresponding control models (Supplementary Tables 2C–2H). Multiple metabolic subsystems, including amino acid metabolism (Figure 5A), vitamin metabolism (Figure 5B), nucleotide metabolism (Figure 5C), fatty acid metabolism (Figure 5D), transport to cellular organelles (Figure 5E), as well as glycolysis and selected TCA cycle reactions (Figure 5F), exhibited significantly higher flux rates in SARS-CoV-, SARS-CoV-2-, and MERS-CoV-infected models compared with controls at both time points (FDR < 0.05; Figure 5; Supplementary Tables 2I and 2J). Increased glycolysis, TCA cycle perturbations, and fatty acid and cholesterol accumulation have been extensively documented in COVID-19 patients,^71–74^ as well as in animal and cellular models of SARS-CoV-2 infection.^40,75–77^ We therefore focused on identifying common and distinct mitochondrial carrier reactions affected by SARS-CoV, SARS-CoV-2, and MERS-CoV.

**Figure 5.**
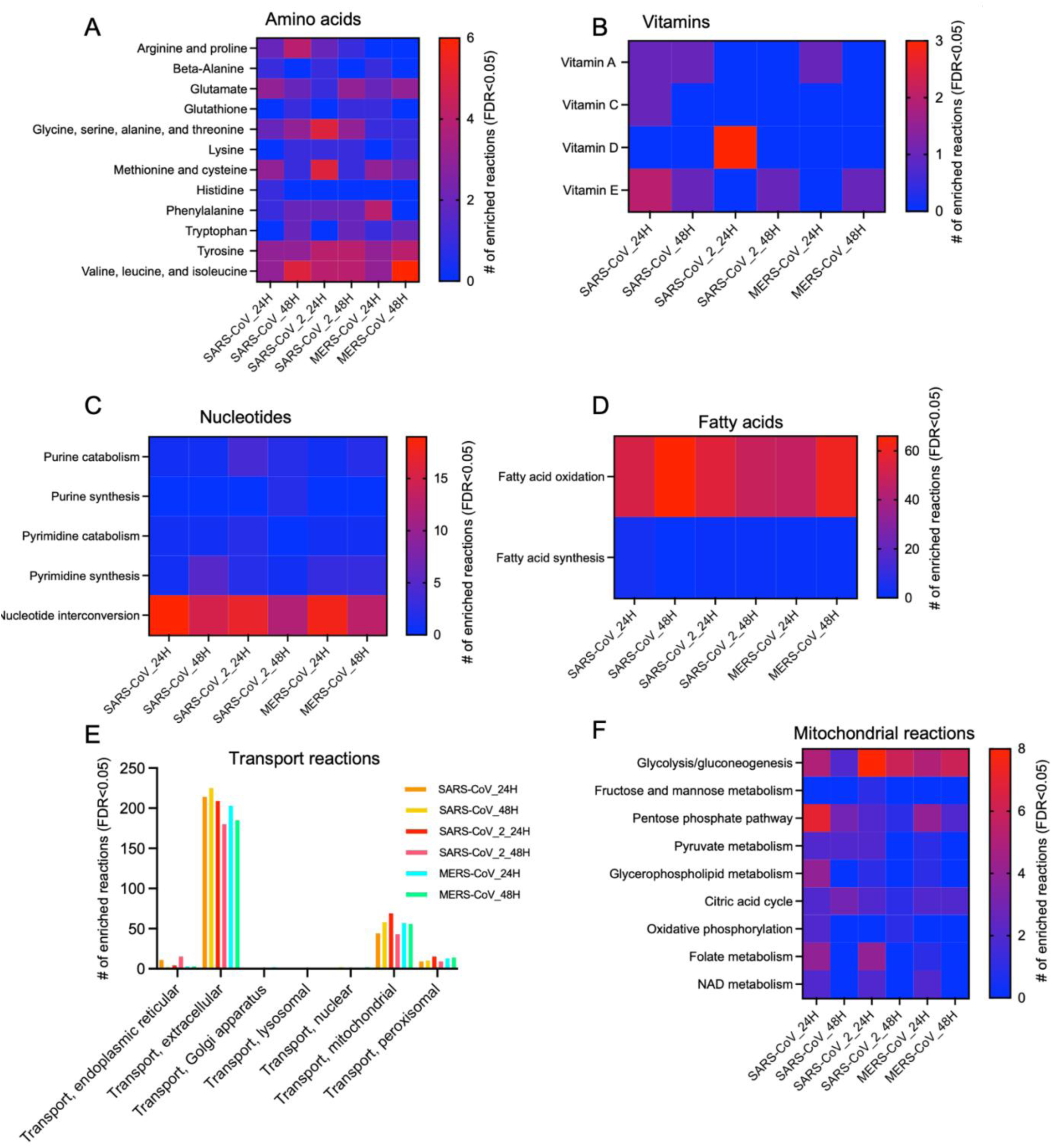
Significantly enriched metabolic subsystems. **(A)**. Enriched amino acids subsystems following infection by SARS-CoV, SARS-CoV-2 and MERS at 24H and 48H post-infection.**(B) (C) (D) (E) and (F)**. As in (A) but vitamins, nucleotides, fatty acids, mitochondrial and transport reactions. Related to **Figure S3**.

Metabolite transport across mitochondrial membranes is mediated by members of the solute carrier family 25 (SLC25), which import and export diverse metabolites, cofactors, and ions.^78^ Notably, expression of 21 of the 53 annotated SLC25 family members was significantly altered following coronavirus infection, with distinct temporal dynamics and directionality across the three viruses (Figure S3). For instance, the aspartate–glutamate carrier SLC25A13 was upregulated by all three viruses at both 24 and 48 hours post-infection, except for MERS-CoV at 24 hours (Figure S3), supporting previous suggestions that SLC25A13 may represent a therapeutic target.^79^ In addition, prolonged coronavirus infection induced upregulation of the ornithine carrier SLC25A15 and downregulation of SLC25A44, a branched-chain amino acid transporter previously implicated in obesity and cardiac pathologies^80^ (Figure S3).

Our analysis identified specific mitochondrial reactions that were commonly affected by all three viruses (Figure 6A), as well as reactions uniquely perturbed by individual coronaviruses (Figure 6C). These alterations highlight deregulation of mitochondrial transport reactions that we classified as “pan-CoV,” shared across all three infections, including reactions involving the carnitine–acylcarnitine carrier (CAC) (Figure 6B). Virus-specific perturbations were also observed (Figure 6B). In particular, nucleotide transport reactions were altered in SARS-CoV-2 and MERS-CoV infections but not in SARS-CoV infection, whereas reactions mediated by the dicarboxylate carrier SLC25A10 (DIC), which exports malate from the mitochondrial matrix in exchange for phosphate, were affected in SARS-CoV and SARS-CoV-2 but not in MERS-CoV (Figure 6B). Collectively, these findings indicate that specific mitochondrial carrier enzymes and transport reactions highlighted here (Figure 6) may represent promising targets for mitigating coronavirus-induced metabolic dysregulation.

**Figure 6.**
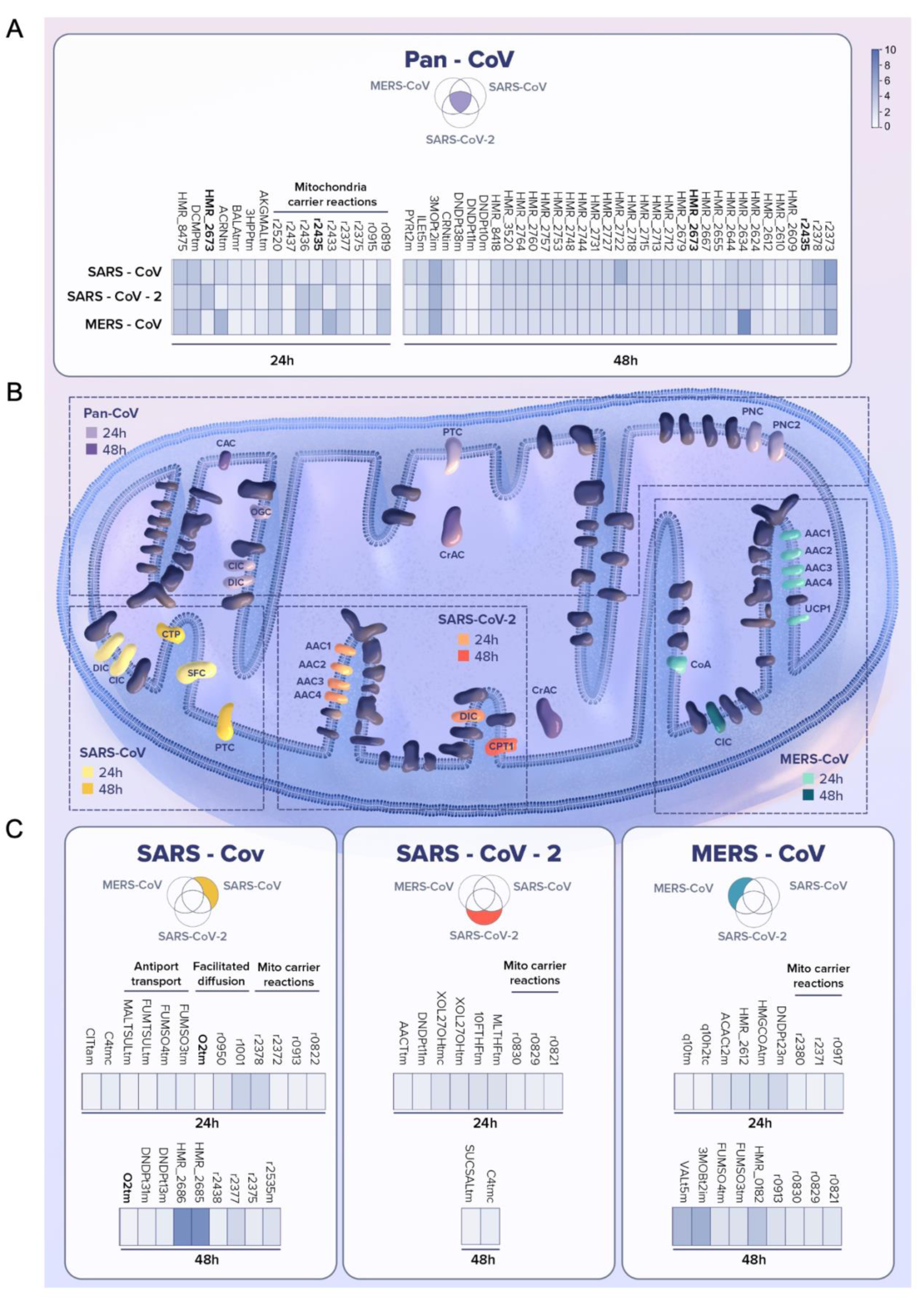
Coronaviruses perturb mitochondria transport reactions. **(A)**. Reactions commonly perturbed by SARS-CoV, SARS-CoV-2 and MERS in the Mitochondrial transport pathway. **(B)**. Schematic representation mitochondria carriers affected by all three coronaviruses (pan-Cov), or specifically by SARS-CoV, SARS-CoV-2 and MERS. **(C)**. As in (A) but reactions specifically perturbed by each indicated virus. See also **Figure S2**.

### An algorithm to predict the rescue of metabolic perturbations

To precisely identify and compare metabolic perturbations in host cells following infection with SARS-CoV, SARS-CoV-2, and MERS-CoV, we developed a methodology termed NiTRO (Network-based integrated Tool for Repurposing Optimization) (Figure S4). NiTRO enables (i) detection of viral-induced metabolic changes, (ii) prediction of the consequences of reaction-flux alterations following in silico double-gene deletions, and (iii) identification of novel metabolic drug-target pairs for each virus of interest. This approach yields a panel of metabolic enzymes perturbed during viral infection, some of which are already targeted in metabolic diseases, thereby creating opportunities for repurposing existing FDA-approved drugs.

This approach extends the metabolic transformation framework introduced by Yizhak et al. (2013)^53^ by evaluating combinatorial (double-gene) rather than single-gene perturbations, and by applying it to the comparative analysis of multiple viral infections. NiTRO integrates genome-scale metabolic models (Recon3D)^52^ with gene inactivity moderated by metabolism and expression (GIMME)^63^ to incorporate RNA-seq expression data from control and infected cells and generate context-specific metabolic models (Figure S4A). Using the Jaccard similarity index, infected models are compared with their corresponding controls to identify a set of “sick reactions,” defined as perturbed reactions in infected models whose flux ranges cover less than 50% of those observed in control models (Figure S4B). In the next step, NiTRO predicts gene knockouts capable of reverting the fluxes of sick reactions toward control-state values (Figure S4C). Specifically, NiTRO computes reaction fluxes following the deletion of every possible gene pair in the infected model, one pair at a time. Using partial least squares dimensional reduction, the algorithm then selects the top 1% of gene-pair deletions, hereafter referred to as rescuer knockouts (KOs), that yield flux profiles linearly correlated with those of the control model (Figure S4C).

Finally, NiTRO identifies reverted reactions by assessing the balance between sick and normal reactions following each rescuer KO: reactions are considered reverted when their post-KO flux ranges cover more than 50% of the corresponding control-model ranges (Figure S4D; Supplementary Tables 3A and 3B). To evaluate NiTRO performance, we quantified the deviation of perturbed reaction fluxes from control values before (Figure 7A–B, grey lines) and after application of the predicted knockouts (Figure 7A–B, blue lines; Supplementary Tables 3C–3H). For a substantial subset of reactions, flux deviations from the control state were reduced following the predicted knockouts. These results demonstrate that rescuer knockouts can partially restore perturbed reaction fluxes toward healthy control states, supporting the performance of the NiTRO algorithm and its potential to predict strategies for reversing infection-induced metabolic perturbations.

**Figure 7.**
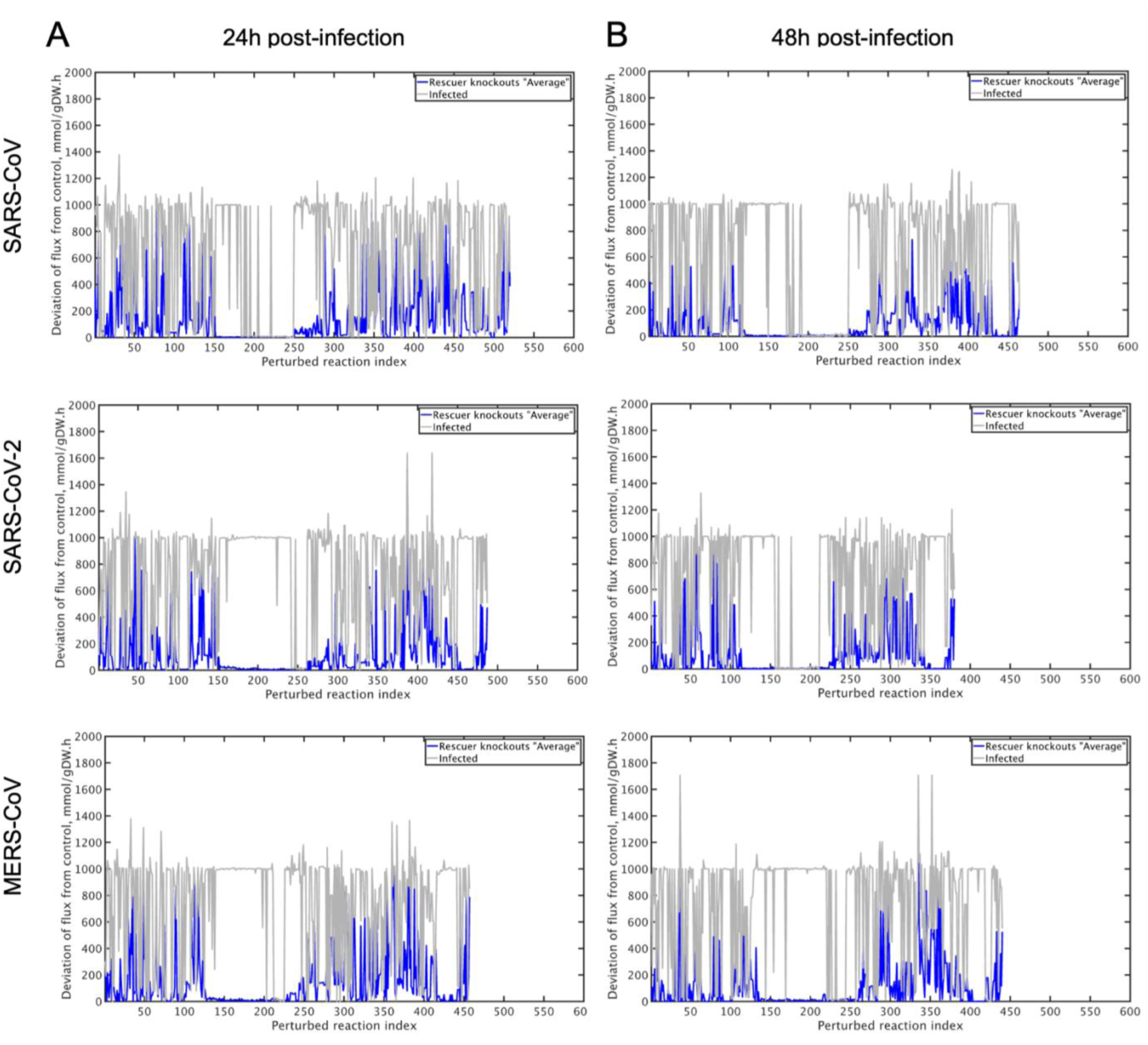
Performance of NiTRO algorithm. Absolute deviation of perturbed reaction fluxes from control-model values, before and after application of NiTRO-predicted rescuer knockouts. (A) 24 hours and (B) 48 hours post-infection. Within each panel, rows correspond to SARS-CoV (top), SARS-CoV-2 (middle), and MERS-CoV (bottom). Each vertical line represents a single perturbed reaction; the x-axis shows the reaction index and the y-axis shows the absolute flux deviation from the corresponding control model (mmol/gDW·h). Grey lines indicate the deviation of each reaction in the infected model prior to any perturbation. Blue lines indicate the average deviation after application of the predicted rescuer gene-pair knockouts. A reduction in line height from grey to blue reflects successful partial restoration of reaction fluxes toward control-model values. Numerical values for each reaction, including control flux, infected-model flux, average rescuer-knockout flux, and corresponding deviations, are provided in Supplementary Tables 3C–3H. Related to Figure S4.

### Predicting sick metabolic reactions following coronavirus infection

To identify perturbed reactions following infection of human cells with SARS-CoV, SARS-CoV-2, and MERS-CoV, we performed flux variability analysis and calculated the flux range of each reaction required to sustain optimal biomass growth. Using the Jaccard index, we compared the flux range of each reaction in infected models with that of the corresponding control model at each time point. We defined “sick reactions” as reactions whose flux ranges in infected models covered less than 50% of the corresponding control-model ranges. Conversely, “healthy” or nonperturbed reactions were defined as those for which infected-model flux ranges covered more than 50% of control-model ranges. Most metabolic reactions and their associated metabolites were not perturbed by coronavirus infection at either 24 or 48 hours post-infection (Figure S5). However, the fluxes of several metabolites displayed patterns that were either common to, or specific for, individual coronaviruses and were associated with distinct sets of sick reactions (Figures 8A–D) at 24 and 48 hours post-infection, respectively. To assess the statistical significance of these observations, we performed random sampling with 10,000 iterations (Figures 8E and 8F). For each iteration, the number of intersecting sick reactions across all possible combinations of infected models was calculated. The observed numbers of sick reactions at 24 hours (Figure 8E, red dots) and 48 hours post-infection (Figure 8F, red dots) were significantly higher than expected from random distributions (Figures 8E and 8F, boxplots). These results indicate that our approach reliably identifies both common and virus-specific sick metabolic reactions following coronavirus infection. Notably, shared sick reactions mapped to metabolic pathways such as nucleotide and carbohydrate metabolism, with subsets appearing preferentially at either 24 or 48 hours post-infection (Figure 8G, red and blue lines).

**Figure 8.**
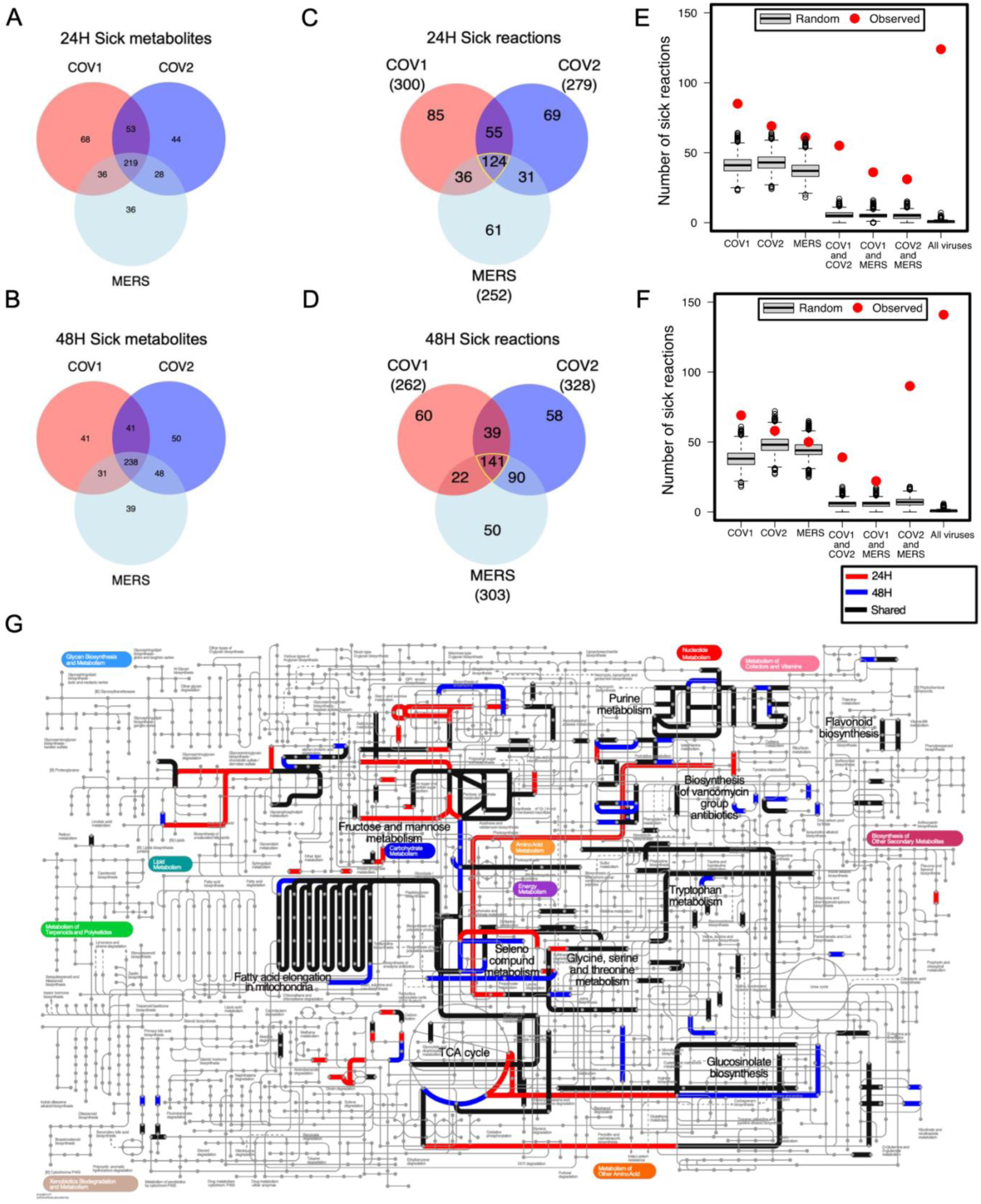
Healthy and sick reactions and associated metabolic pathways. **(A-D)** We use the term “sick metabolite or reaction” when the flux range of the metabolite or reaction in the infected model covers less than 50% of that in the control models. The Venn diagrams show the number of sick metabolites (**A** and **B**) or sick reactions (**C** and **D**) that are common or exclusive among the infected models at 24H and 48H post-infection. (**E** and **F**) Observed distributions compared to random sampling at 24H (E) and 48H (F) post-infection. (**G**) iPath3 visualization of the pathways that are assigned to the subset of sick reactions at each timepoint. Red and blue lines correspond to sick reactions that are exclusively found at 24H, or 48H, respectively. Black lines, correspond to sick reactions that are common between both timepoints.

### Druggable rescuer metabolic gene pairs following coronavirus infection

To identify druggable metabolic gene pairs capable of rescuing infection-induced metabolic perturbations, we applied the NiTRO algorithm (Figure S4) to compute gene pairs participating in sick reactions that, when deleted, yielded reaction flux profiles in infected models that correlated linearly with those of control models. We identified multiple gene-pair deletions predicted to rescue sick reactions following infection with SARS-CoV, SARS-CoV-2, MERS-CoV, or all three viruses at both 24 and 48 hours post-infection. We next filtered these gene pairs to retain only those in which both genes are targets of existing FDA-approved drugs and visualized the resulting networks using Cytoscape^81^ (Figure S6; Supplementary Table 3I). This analysis revealed that genes encoding SLC5A3, IDH1, and IDH2 act as druggable hubs, with potential relevance for SARS-CoV-1 and SARS-CoV-2 infection. Our analysis further identified key double-knockout (KO) gene pairs capable of shifting infected cells from a virus-induced metabolic state toward a healthy metabolic condition. An overview of these KOs is presented using Venn diagrams showing shared and virus-specific KOs across SARS-CoV, SARS-CoV-2, and MERS-CoV at 24 and 48 hours post-infection (Figure 9A,B), together with gene interaction networks illustrating KO interactions at the same time points (Figure 9C–H). At 24 hours post-infection, 3,145, 3,063, and 749 KO gene pairs were specific to SARS-CoV, SARS-CoV-2, and MERS-CoV, respectively, with 707 KO pairs shared among all three viruses (Figure 9A). These shared KOs suggest common rescue mechanisms targeting core metabolic pathways essential for viral replication. By 48 hours post-infection (Figure 9B), virus-specific dependencies became more pronounced, as evidenced by increased numbers of double-KO gene pairs required to rescue infected phenotypes. SARS-CoV-2 exhibited the highest number of unique KOs at 48 hours (n = 4,592), followed by SARS-CoV (n = 3,159) and MERS-CoV (n = 2,936). Gene interaction network analysis (Figure 9C–H) further highlighted virus-specific metabolic dependencies by revealing central hub genes within the KO networks. At 24 hours, hub genes such as FH, CPT2, and CAT were prominent in SARS-CoV and SARS-CoV-2 networks, whereas fewer dominant hubs were observed for MERS-CoV. By 48 hours, the network topology shifted, with GLB1, SDHA, and DTYM emerging as key hub genes, reflecting an increasing reliance of the viruses on specific metabolic pathways as infection progresses.

**Figure 9.**
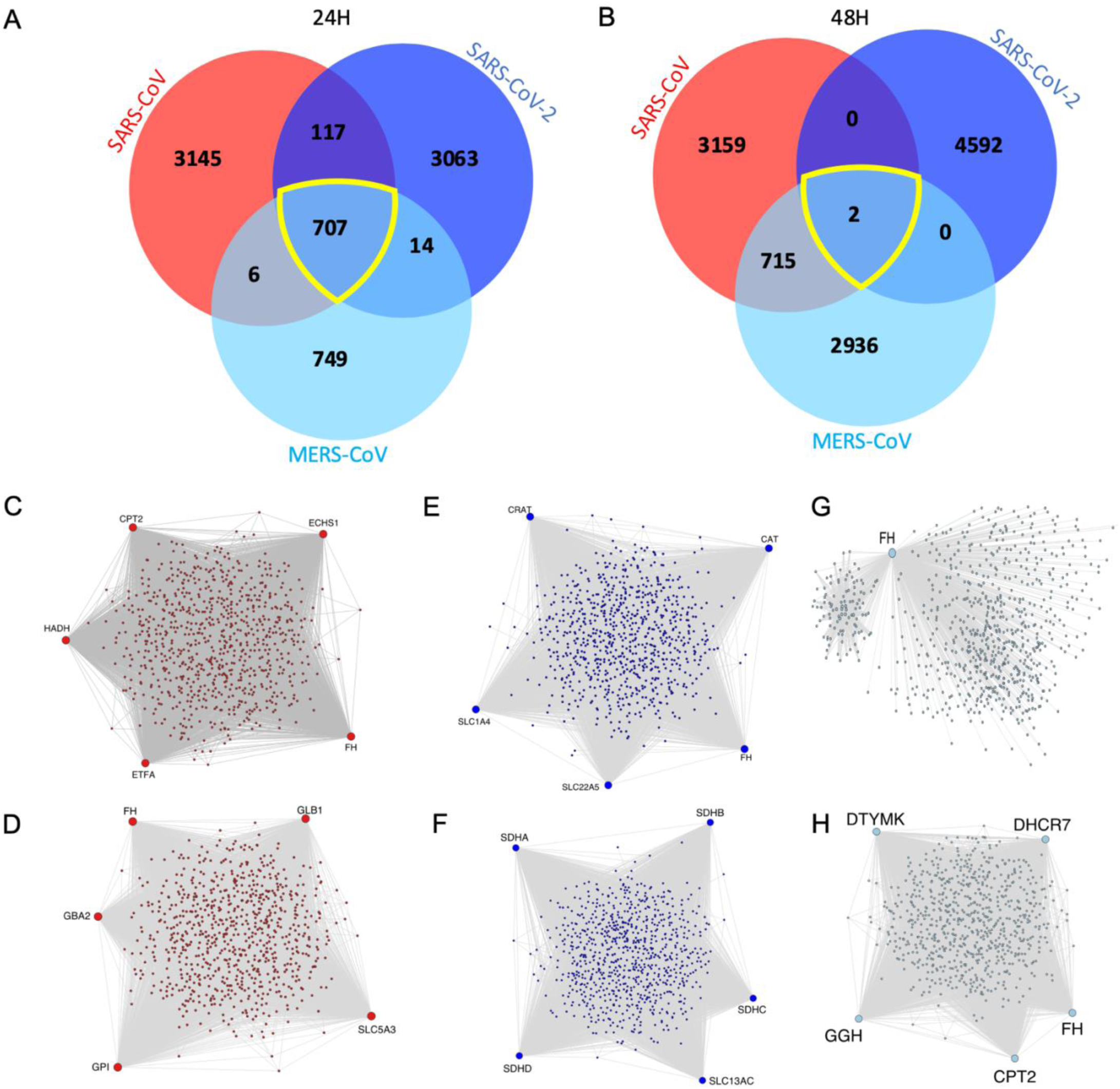
Rescuer knockouts and networks highlighting key hub genes. (**B**) Venn diagrams representing rescuer knockout (KO) counts identified at 24 hours and 48 hours post-infection by indicated viruses, respectively. Overlaps show shared rescuer KOs. **(C-H)** depict gene interaction networks where nodes represent genes and edges link gene pairs that correspond to rescuer knockout (KO) interactions. The colors of the nodes and edges indicate the virus from which the KOs were identified: red for SARS-CoV, blue for SARS-CoV-2, and light blue for MERS-CoV. Panels **(C)**, **(E)** and **(G)** show interactions at 24 hours post-infection.

Notably, while 707 rescuer KO pairs were shared among all three viruses at 24 hours post-infection (Figure 9A), this shared signal collapsed almost entirely by 48 hours, with zero shared KOs between SARS-CoV and SARS-CoV-2, zero between SARS-CoV-2 and MERS-CoV, and only 2 between SARS-CoV and MERS-CoV (Figure 9B). This temporal divergence indicates that, whereas early infection engages conserved host metabolic dependencies amenable to broad-spectrum intervention, prolonged infection drives virus-specific metabolic rewiring that necessitates pathogen-tailored targeting strategies.

### Concordance of NiTRO predictions with independent experimental and clinical evidence

To assess whether NiTRO identifies biologically meaningful rescue targets, we asked whether its top-ranked predictions correspond to genes and pathways for which independent experimental or clinical evidence of antiviral efficacy already exists. Concordance between NiTRO predictions and independently obtained results serves to validate the computational framework rather than to claim novelty of individual drug– target associations.

Several highly ranked NiTRO predictions map directly onto genes with independently validated roles in SARS-CoV-2 infection (Table 1). Metformin, predicted by NiTRO to reverse CPT2- and SDHA-mediated flux perturbations via complex I inhibition, was shown to inhibit infectious SARS-CoV-2 by up to 99% in Calu-3 and Caco-2 cells.^82^ In the phase 3 COVID-OUT trial, early metformin treatment reduced the incidence of Long-COVID or death by ∼41% at six months,^83^ and a recent meta-analysis of ≈ 2.9 million diabetic patients reported a 22% reduction in COVID-19 mortality (adjusted OR 0.78, 95% CI 0.69–0.88; *p* < 0.00001).^84^

**Table 1.**
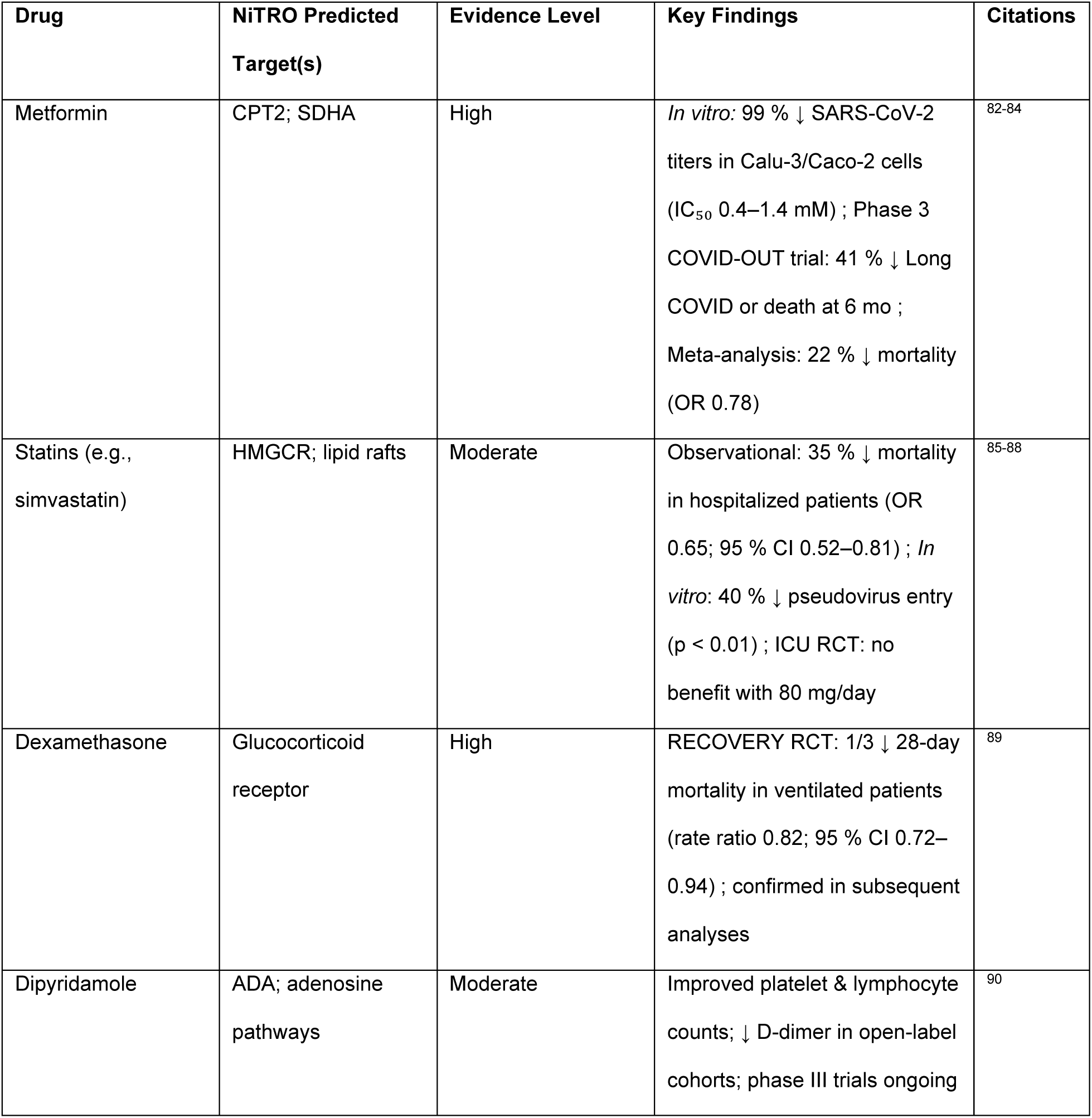
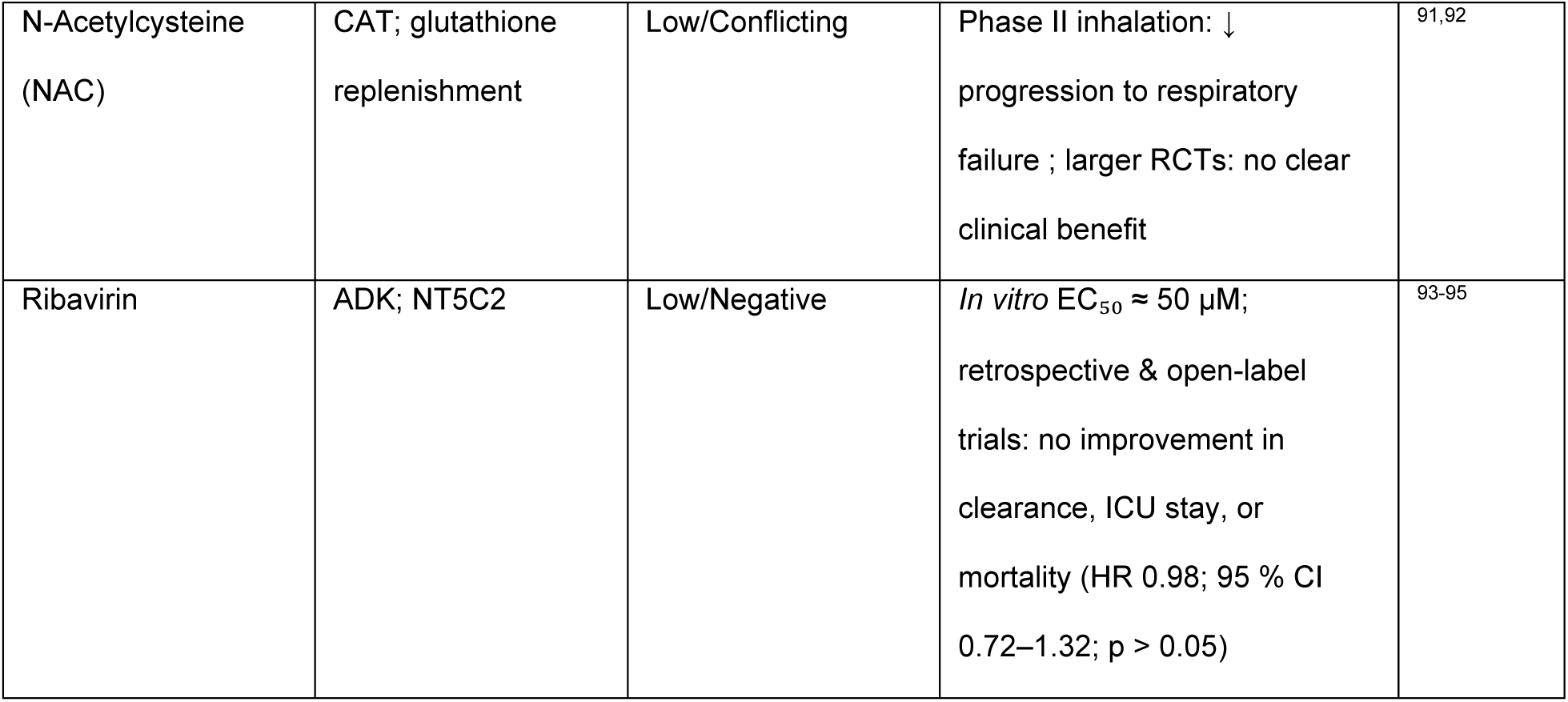
Concordance of NiTRO predictions with independent experimental and clinical evidence. *Drug names, NiTRO predicted targets, evidence level, key findings and citations are shown*.

Similarly, statins targeting HMGCR and disrupting lipid-raft–mediated viral entry, were associated with a 35% lower mortality in hospitalized cohorts (OR 0.65, 95% CI 0.52– 0.81)^85,86^ and reduced SARS-CoV-2 pseudo-virus entry by ∼40% *in vitro* (*p* < 0.01),^87^ although a large adaptive RCT in critically ill patients found no benefit of high-dose simvastatin on days alive without organ support ^88^. Dexamethasone, while not a direct NiTRO hit, illustrates the value of host-directed repurposing: low-dose regimens cut 28-day mortality by one-third in ventilated patients (rate ratio 0.82, 95% CI 0.72–0.94) and are now standard of care for severe COVID-19.^89^ Dipyridamole, implicated via ADA, was shown to improve platelet and lymphocyte counts and lowered D-dimer levels in small open-label cohorts,^90^ with phase III trials underway. N-acetylcysteine (NAC), intended to bolster CAT-mediated redox balance, reduced progression to respiratory failure in a phase II inhalation trial,^91^ but failed to show clear benefit in larger RCTs.^92^

By contrast, ribavirin, linked to ADK and NT5C2 flux modulation, despite an EC₅₀ ≈ 50 µM in Vero E6 cells, did not improve viral clearance, ICU stay, or mortality in clinical studies (adjusted HR 0.98, 95% CI 0.72–1.32; p > 0.05).^93^ Other predicted rescuers such as IDH1/IDH2, SDHA, FH, and CAT remain at the preclinical stage, supported only by preliminary in cellular or animal data (e.g., SIRT5-mediated desuccinylation reducing replication by ∼50% in A549 cells;^94^ IDH1 inhibition lowering lung titers by 1.5 log₁₀ without survival benefit^95^ and no registered COVID-19 trials to date.

In sum, NiTRO’s top predictions, metformin and statins, have the most robust multi-level support (in cellulo, clinical trials, and large observational cohorts). Dipyridamole and NAC show mixed but promising early signals, whereas ribavirin has been refuted clinically. Remaining hits (IDH1/IDH2, SDHA, FH, CAT) lack clinical validation and warrant focused preclinical exploration (See Table 1 and Figure S7).

## DISCUSSION

The results presented here establish NiTRO as a genome-scale computational framework capable of translating virus-induced transcriptional reprogramming into ranked, mechanistically grounded therapeutic hypotheses. Three features of the data collectively define the paper’s contribution: first, the compression of extensive transcriptional divergence into a constrained set of convergent metabolic perturbations; second, the sharp temporal bifurcation in shared metabolic vulnerabilities between 24 and 48 hours post-infection; and third, the alignment between the highest-confidence NiTRO predictions and independently validated clinical interventions. Together, these features argue that genome-scale flux modeling adds a layer of interpretive resolution to transcriptomic data that gene expression analysis alone cannot provide^65^, and that the resulting target nominations carry mechanistic grounding.

### From transcriptional divergence to metabolic convergence

A central and non-obvious finding is that three viruses that collectively dysregulate thousands of host genes, with only 775 and 665 in common across all three at 24 and 48 hours respectively, converge at the flux level on a statistically significant and constrained set of sick reactions enriched in mitochondrial transport, nucleotide biosynthesis, fatty acid metabolism, and branched-chain amino acid catabolism. This compression is not incidental: it reflects the topological architecture of metabolic networks, in which distributed transcriptional perturbations funnel through a limited number of stoichiometric and thermodynamic bottlenecks.^96^ The GIMME-based contextualization^63^ and subsequent flux variability analysis perform precisely this filtering step, and it is what makes the downstream NiTRO target nominations more actionable than pathway enrichment analysis of differentially expressed genes. The sick reactions are not associations; they are flux-level consequences of the observed transcriptional state, constrained by the biochemical logic of Recon3D.^52^ The simultaneous activation of nucleotide processing, folate, and amino sugar metabolism alongside deactivation of fatty acid oxidation, glycolysis, and steroid catabolism at 24 hours is consistent with a coordinated anabolic reorientation, which is a shift from catabolic energy extraction toward biosynthetic precursor supply that mirrors the Warburg-like metabolic reprogramming documented in SARS-CoV-2 infection and in other viral systems.^40,97^ Notably, this activation-deactivation pattern at 24 hours is shared across all three viruses despite their distinct transcriptional signatures, reinforcing that the metabolic layer imposes a convergent logic on divergent upstream gene expression. By 48 hours, this shared anabolic program begins to fragment: nucleotide metabolism equilibrates, cholesterol metabolism and tryptophan catabolism are activated, and oxidative phosphorylation and redox balance pathways are deactivated — a transition more consistent with accumulating bioenergetic stress than with continued productive exploitation of host biosynthetic capacity. The directionality reversal of mitochondrial pyruvate kinase and pyruvate phosphotransferase reactions at 48 hours, detected through the flux sign analysis, is particularly informative in this regard, indicating altered equilibria in central carbon metabolism that are not captured by fold-change analysis of transcript levels alone.^98^

### Mitochondrial carriers as a pan-coronavirus vulnerability class

The perturbation of 21 of 53 annotated SLC25 family members across coronavirus infection, with distinct temporal dynamics and directionality across the three viruses, establishes the mitochondrial transporter system as a qualitatively different category of viral target than the enzymatic hubs identified by NiTRO. Enzymatic hubs can in principle be circumvented by metabolic rerouting; transporters that gate substrate entry to the mitochondrial matrix have fewer bypass routes.^78,99^ The universal upregulation of SLC25A13 (aspartate-glutamate carrier) across all three viruses at 48 hours, and by SARS-CoV and SARS-CoV-2 at 24 hours, is particularly notable: this transporter drives the malate-aspartate shuttle linking cytosolic and mitochondrial redox states, and its consistent targeting suggests that all three viruses require enhanced aspartate-glutamate exchange to sustain the elevated metabolic flux observed globally. The carnitine-acylcarnitine carrier (CAC) emerging as a pan-coronavirus perturbed reaction, which is directly linked to the CPT2-mediated fatty acid import pathway that is a prominent 24-hour rescue hub, makes the SLC25 family a natural target class for broad-spectrum antiviral development, complementing rather than duplicating the enzymatic hub nominations of NiTRO. The virus-specific divergence in SLC25A10 (dicarboxylate carrier, malate-phosphate exchange; altered in SARS-CoV and SARS-CoV-2 but not MERS-CoV) and nucleotide transport (altered in SARS-CoV-2 and MERS-CoV but not SARS-CoV) further demonstrates that the SLC25 family is a site of both convergent and individualized exploitation, with implications for both pan-coronavirus and tailored intervention strategies.

### The therapeutic window defined by temporal bifurcation

The collapse of shared rescuer knockouts from 707 across all three viruses at 24 hours to near-zero overlap by 48 hours, zero between SARS-CoV and SARS-CoV-2, zero between SARS-CoV-2 and MERS-CoV, two between SARS-CoV and MERS-CoV, is the most consequential quantitative finding in the paper for drug strategy. It defines a time-bounded window during which broad-spectrum host-directed metabolic intervention is mechanistically feasible, and after which each virus has established sufficiently distinct metabolic dependencies that any effective intervention must be pathogen-tailored. The hub gene transition underlying this collapse is interpretable: at 24 hours, FH (TCA cycle fumarate hydration), CPT2 (mitochondrial fatty acid import), and CAT (hydrogen peroxide detoxification) are prominent in SARS-CoV and SARS-CoV-2 rescue networks, representing the metabolic infrastructure for the initial burst of replication — energy, biosynthetic carbon, and oxidative stress management. By 48 hours, GLB1, SDHA, and DTYM dominate, pointing toward lysosomal processing, sustained TCA cycle-to-electron transport chain coupling, and thymidylate synthesis for ongoing nucleotide supply. This succession is not random: it suggests viruses first secure the basic energy supply and then progressively co-opt the specialized biosynthetic programs^66^ required for sustained virion production, an escalating and deepening metabolic siege whose early phase is shared and whose later phase is individualized. This temporal architecture provides a systems-level, flux-biology rationale for a consistently observed clinical phenomenon: antiviral interventions are most effective when initiated early.^100^ The usual explanation invokes lower viral burden at early time points.^101,102^ The NiTRO data suggest an additional, mechanistically distinct explanation: the shared metabolic vulnerabilities that make broad-spectrum host-directed intervention feasible are only accessible during a narrow early window that closes as virus-specific rewiring matures. Patients who present to care after the 48-hour window has closed are not simply at a higher viral burden; they are in a metabolically distinct disease phase where the pan-coronavirus rescue targets are no longer rate-limiting. This reframes the clinical urgency of early treatment in metabolic terms and has design implications for trials of host-directed metabolic interventions: enrollment criteria should capture early presenters, and time-from-symptom-onset should be treated as a primary stratification variable rather than an enrollment filter.

### Mechanistic interpretation of the clinical validation gradient

The concordance analysis in Table 1 functions as external validation of the NiTRO framework, and the gradient of clinical support across the nominated targets is itself mechanistically interpretable. Metformin achieves the highest validation tier, 99% *in vitro* inhibition of SARS-CoV-2 in Calu-3 and Caco-2 cells,^82^ 41% reduction in Long-COVID or death in the Phase 3 COVID-OUT trial,^83^ and 22% mortality reduction across approximately 2.9 million patients in meta-analysis.^84^ The mechanistic basis for this exceptional performance is explicable from the NiTRO results: metformin inhibits mitochondrial complex I, which perturbates both CPT2-dependent fatty acid oxidation (the dominant 24-hour rescue hub) and SDHA-linked TCA cycle function (a dominant 48-hour hub).^103^ This dual temporal coverage, addressing both early and late infection-phase vulnerabilities through a single compound’s mechanism of action, distinguishes metformin from single-node interventions and likely accounts for its breadth of observed benefit. The Long-COVID efficacy in particular is consistent with metformin’s sustained activity against the metabolic states that persist beyond acute infection,^104,105^ a pattern that single-target interventions addressing only the acute-phase hubs would not be expected to replicate. Statins achieve their benefit through a mechanistically distinct route — HMGCR inhibition disrupting cholesterol synthesis and lipid-raft-mediated viral entry^106^ — that is parallel to, rather than downstream of, the core NiTRO rescue hubs. Their clinical signal (35% lower mortality in observational cohorts^107^; 40% pseudovirus entry reduction *in vitro*^108^) is genuine but the negative result in the REMAP-CAP adaptive RCT in critically ill patients^88^ deserves interpretation rather than dismissal: the trial enrolled patients at ICU admission, almost certainly after the 24-hour convergence window had closed and in a disease context dominated by immunopathology rather than active viral replication. The statin mechanism acts on viral entry, which is most relevant before sustained replication is established, which is precisely the phase the trial systematically excluded. The observational evidence, which includes earlier-stage hospitalized patients, more likely captures the window during which the mechanism is actionable. NAC’s failure in larger RCTs^92^ despite targeting CAT-mediated redox balance, a genuine 24-hour hub, illustrates a principle about hub hierarchy that the NiTRO data make visible: not all hubs in the rescue network are equally rate-limiting for viral replication. CAT occupies the redox subsystem of the rescue network, and its perturbation restores reactive oxygen species balance without addressing the upstream CPT2 and FH perturbations that drive the broader metabolic rewiring. Restoring a downstream consequence while leaving upstream causes intact is unlikely to produce durable benefit. Ribavirin’s clinical failure (HR 0.98),^93^ despite targeting ADK and NT5C2 in the nucleotide metabolism subsystem that is genuinely perturbed, likely reflects that nucleotide pool depletion is not the rate-limiting step for viral replication in this context — the viral genome is being replicated within a cell that NiTRO shows has activated nucleotide biosynthesis, making nucleotide supply less of a bottleneck than its perturbation in isolation would suggest. Together, these cases suggest that the clinical translatability of NiTRO-nominated targets correlates with whether the target sits on a rate-limiting flux route to viral replication, not merely whether it belongs to a perturbed pathway.

### Druggable hubs and the preclinical frontier

The identification of SLC5A3, IDH1, and IDH2 as druggable hub genes connecting multiple rescuer knockout pairs in the NiTRO network positions them as priority candidates for focused experimental validation. IDH1 and IDH2 generate alpha-ketoglutarate and NADPH, metabolites that serve dual and competing roles in viral replication and host antioxidant defense, creating a genuine metabolic conflict point that is distinct in character from the energy supply hubs CPT2 and SDHA. Preliminary preclinical evidence is consistent with the nomination: IDH1 inhibition reduced lung viral titers by 1.5 log₁₀ in an animal model,^95^ and SIRT5-mediated desuccinylation of SDHA reduced replication by approximately 50% in A549 cells.^94^ Neither has yet reached clinical validation in a coronavirus context, and both warrant systematic investigation in physiologically relevant infection models before their position in the therapeutic hierarchy can be established. SLC5A3, as an inositol transporter, connects the NiTRO rescue network to phosphoinositide signaling and membrane remodeling.^109^ This metabolic territory has received limited attention in the antiviral context but that represents an interesting mechanistic bridge between the metabolic and membrane biology of coronavirus infection.

### Limitations and generalizability

The interpretation of these results should account for several methodological constraints. The exclusive use of HEK293-ACE2 cells was a deliberate design choice — minimizing confounding from tissue-specific differentiation programs and heterogeneous receptor expression to isolate core metabolic responses, but it necessarily limits inference to the metabolic responses that are cell-type-independent. Primary airway epithelial cells, alveolar type II pneumocytes, and macrophages each carry distinct metabolic programs that may alter both the perturbation landscape and the rescuer hierarchy.^110^ The GIMME algorithm’s binary gene activity threshold suppresses expression gradients that may carry metabolically relevant information; algorithms that preserve continuous expression variation in model contextualization may identify additional or different flux perturbations.^111^ FBA at steady state cannot resolve the rapid transient flux states^65^ of early viral entry and the first wave of replication; the 24-hour models capture a metabolic snapshot that integrates events between 0 and 24 hours rather than resolving their sequence. And the two-timepoint design, while sufficient to demonstrate temporal divergence, cannot resolve the kinetics of hub transition or identify vulnerabilities that are transiently prominent and then resolved. Finer temporal resolution at six, 12, 18, 24, 36, and 48 hours would reveal whether the convergence-to-divergence transition is abrupt or gradual, and whether additional therapeutic windows exist within the early infection phase.

The NiTRO framework is, in principle, applicable to any pathogen for which matched infected and uninfected transcriptomic data exist in a cell type covered by an available genome-scale metabolic reconstruction. The comparative strategy applied here — identifying the temporal intersection of pathogen-specific metabolic perturbation landscapes to nominate conserved, host-encoded targets — offers a resistance-robust complement to direct-acting antiviral approaches: host metabolic enzymes cannot accumulate escape mutations under viral selection pressure.^112^ As transcriptomic datasets from diverse viral infections expand and metabolic reconstructions become available for additional cell types and tissues, the NiTRO strategy scales naturally toward a pan-pathogen therapeutic target identification platform. The validation presented here — most compellingly the metformin Phase 3 trial result — establishes proof of concept that this approach can recover clinically actionable targets from transcriptomic data alone, transforming genome-scale metabolic modeling from a descriptive systems biology tool into a prospective component of the antiviral drug discovery pipeline.

## STAR METHODS

### Resource availability

#### Lead contact

Further information and requests for resources and reagents should be directed to and will be fulfilled by the lead contact, Kourosh Salehi-Ashtiani.

#### Materials availability

This study did not generate new unique reagents.

#### Data and code availability

- All data used to construct the models are available at 10.5281/zenodo.17054054.
- All sequencing data are available at the National Center for Biotechnology Information (NCBI) in BioProject ID PRJNA1279424.
- All original code has been deposited at the above Zenodo repository and is publicly available as of the date of publication.
- Any additional information required to reanalyze the data reported in this paper is available from the lead contact upon request.

### Experimental model and study participant details

#### Cell lines

Human embryonic kidney cells (HEK293-ACE2) were cultured in Dulbecco’s Modified Eagle medium (DMEM, Sigma-Aldrich, St. Louis, MO, USA; Cat# D5796) containing 10% of fetal bovine serum (FBS, Sigma-Aldrich; Cat# F7524), 2 mM of L-glutamine (Sigma-Aldrich; Cat# G7513), 100 units of penicillin and 100 µg/mL of streptomycin (Sigma-Aldrich; Cat# P4333) in a cell culture incubator at 37°C with 5% CO2 and 95% humidity.

#### Viruses

SARS-CoV-1, SARS-CoV-2, and MERS-CoV were used at a multiplicity of infection (MOI) of 0.01. The reference sequences used are: human genome GRCh38, viral reference sequences NC_004718.3 for SARS-CoV-1, NC_019843.3 for MERS-CoV, and NC_045512.2 for SARS-CoV-2.

### Method details

#### RNA extraction and sequencing

Twenty-four hours after seeding, HEK293-ACE2 cells (*n* = 3 per group) were infected or not with SARS-CoV-1, SARS-CoV-2, or MERS-CoV viruses, respectively at a multiplicity of infection (MOI) of 0.01 (*n* = 18 total groups). The medium was replaced with fresh medium after two hours, and infection was stopped at 24H or 48H post-infection. Total RNAs were extracted using Nucleospin RNA extraction kit (Macherey-Nagel, Düren, Germany, Cat# 740955) according to the manufacturer’s instructions, and the integrity of RNA verified using Agilent 2100 Bioanalyzer Systems (Agilent Technologies, Santa Clara, CA, USA). Libraries were prepared using the Illumina Stranded mRNA Ligation kit (Illumina, San Diego, CA, USA; Cat# 20040532), and sequenced with NovaSeq 6000 – 300 cycles standard workflow (Illumina, San Diego, CA, USA). The sequencing quality was evaluated by fastQC version 0.11.9, and sequence reads were processed using Qiagen CLC genomics package.

#### Generating context-specific genome-scale metabolic models

Gene expression mRNA data for HEK293-ACE2 cells before and after 24 and 48 hours infection with SARS-CoV, SARS-CoV-2, and MERS-CoV, were integrated with the COBRA human model, Recon3D (Brunk et al., 2018). The integration step uses GIMME algorithm,^63^ available in the COBRA toolbox.^113^ GIMME requires binary entries for the indication of the absence and presence genes. Therefore, we used the 50^th^ percentile TPM gene expression value as the threshold for a gene to be considered as present. Genes with expression values below the threshold were given a value of 0 (absent), and those with expression values above the threshold were given a value of 1 (present). GIMME only activates reactions associated with present genes, generally leaving those associated with the lowly expressed genes inactive. However, GIMME reactivates the reactions associated with lowly expressed genes if they are required to enable the context-specific metabolic network to grow.

#### Flux balance analysis (FBA)

FBA calculates the flow of metabolites through a metabolic network, thereby predicting the flux of each reaction contributing to an optimized biological objective function c, such as growth rate.^114^ Simulating growth rate is represented by a reaction that corresponds to the conversion of precursor metabolites into biomass components such as lipids, nucleic acids, and proteins. The same general biomass objective function was used for all simulation in this study including FBA, FVA, and double gene deletions. The general biomass reaction is defined in Recon3D.{Brunk, 2018 #33} When performing FBA, the flux solution space is constrained.

For each of the 8 models generated after the integration step, we used the biomass objective function as defined in the Recon3D model to obtain FBA solutions using the COBRA toolbox command, *optimizeCbModel.* After setting the objective function in the model, the entries to the command *optimizeCbModel* are the model and the required optimization of the objective function (‘max’ or ‘min’), which in our studies is the maximization of the biomass production. The minNorm was set to 1e^-6^ to minimize the squared Euclidean Norm of internal fluxes, providing a unique, regularized optimal flux solution.^114^ The command output is the FBA solution, which includes the value of the maximum biomass production rate and a column vector with the reaction fluxes for each reaction accounted for in the model, resulting in a stoichiometric matrix with metabolites and reactions.^114^ The term represents the transposed weight vector *C*, which indicates how much each reaction () contributes to the linear objective function.^114^ The net reaction flux vector is given by the total rate of metabolite production, which equals the total rate of consumption. Furthermore, each reaction rate is constrained within a lower and upper bound, respectively. By convention, in this study, we set infinite bounds on internal reaction rates,^114^ with reversible reactions ranging between -1000 and 1000, and irreversible reactions ranging between 0 and 1000.

#### Identification of perturbed reactions

A reaction with a non-zero flux is considered active, and a reaction with a zero flux is considered inactive. Inactive reactions are unable to contribute in the biomass production. A reaction that is active in the infected model and inactive in the control model or vice versa, is identified to carry a perturbation on its activation status. We identified the reaction subset associated with perturbation on activation status by comparing the flux vector of the control model to that of infected model at the respective timepoint. The reaction subsets are shown in Figure 4. The fold change (FC) of a reaction flux is the ratio of its flux in the infected model to that in the control model. Any reaction with |log₂ FC| > 1 was identified to carry a perturbation on its flux value.

#### Flux variability analysis (FVA)

FVA calculates the minimum and maximum range of each reaction flux when maximal possible biomass production rate is satisfied (https://doi.org/10.1186/1471-2105-11-489). We used *fluxvariability*, a command implemented in the COBRA Toolbox for performing FVA. For each of the 8 models, we used the COBRA toolbox command, *fluxvariability*, to obtain the FVA solution. After setting the objective function in the model, the entries to the command *fluxvariability* are the model and the required optimization of the objective function (‘max’ or ‘min’), which in our studies is the maximization of the biomass production. The command outputs are the minimum and maximum flux range for each reaction when biomass production rate is maximum.

#### Gene pair deletion

Removing a gene from the model deactivates the reactions catalyzed by its encoded enzyme. This removal is implemented in the model by setting the upper and lower boundaries of the reactions that associate the encoding gene to zeros. We performed a pairwise gene deletion by eliminating a gene pair from the model. After each gene pair deletion, we performed an FBA to obtain the flux vector.

We performed a deletion of all possible gene pairs, once at a time, for each of the 6 infected models (∼500 000 gene pairs per model). The output is the knockout matrix which is a matrix with m rows equals to the number of model reactions (∼2400 reactions per model), and n columns equals to the number of gene pairs (n = ∼ 500 000). Each column of this matrix is the flux vector representing the consequence of deleting a gene pair on reaction fluxes.

#### Partial least squares analysis (PLSA)

PLSA is a principal component technique for data with high dimensions. It is used to group variables based on correlation among them. In PLSA, a variable can be selected as the objective for evaluating its correlation with other variables. The loadings on the first component correspond to the linear correlation between the variables and the objective variable. For this study, we considered the flux vector obtained for the control model as the objective variable. We used the model-knockout matrix and the objective flux vector as the inputs to the PLSA. We considered a deleted gene pair yielding to a flux vector with a loading higher than the 99^th^ percentile (top 1%), as a rescuer knockout.

### Quantification and statistical analysis

Statistical tests used throughout this study are described in the figure legends and Method details. Differential gene expression analysis was performed using standard RNA-seq pipelines with FDR correction. The Kolmogorov-Smirnov test was used to compare flux distributions between infected and control models. Random sampling with 10,000 iterations was performed to assess the statistical significance of shared sick reactions across infected models. A reaction was considered perturbed if its flux deviated by at least two-fold (|log₂ FC| ≥ 1) from the control model. Pathway enrichment analyses were performed with FDR < 0.05 as the significance threshold. Rescuer knockouts were defined as gene-pair deletions yielding flux vectors with PLS loadings above the 99th percentile.

### Key resources table

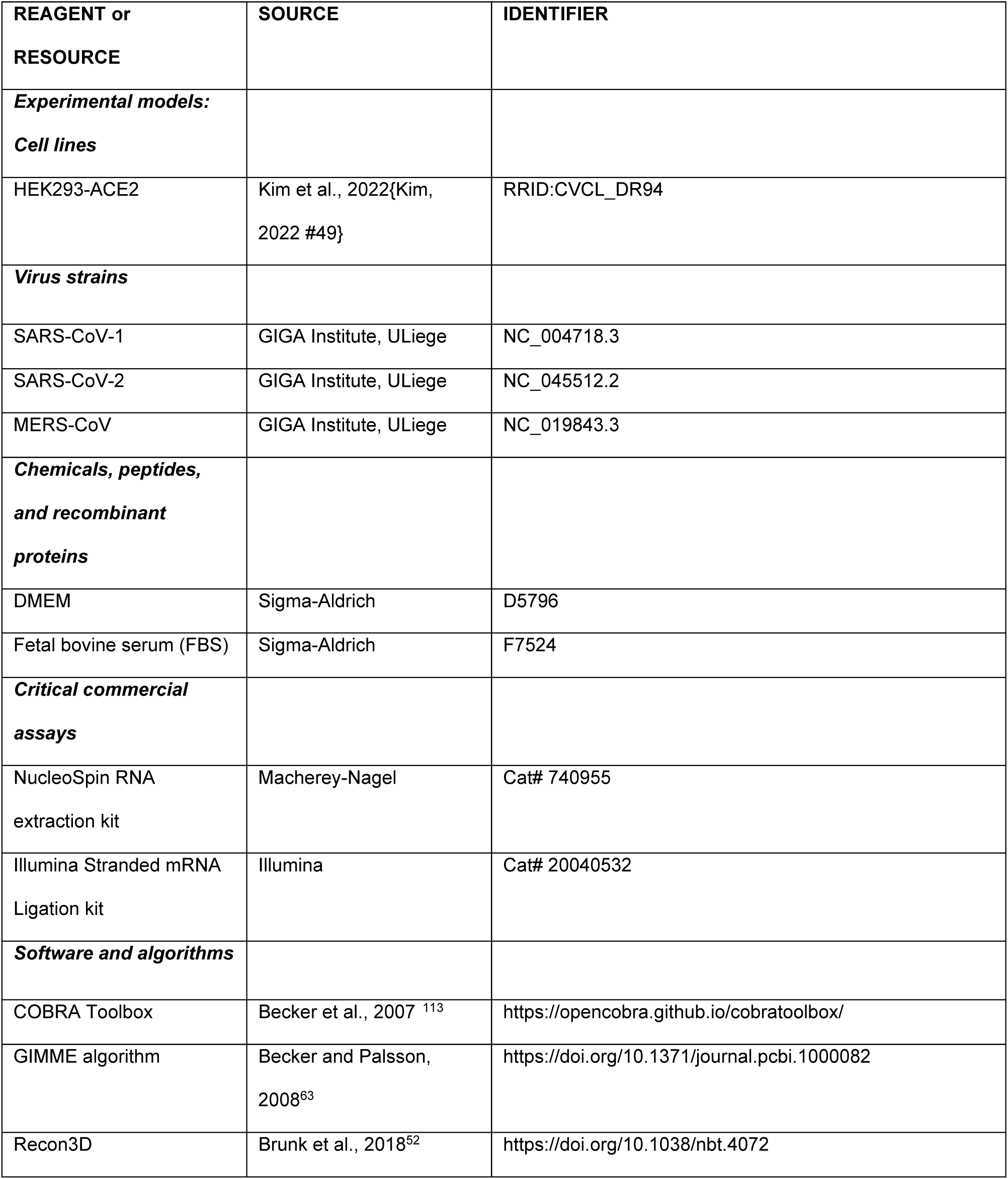
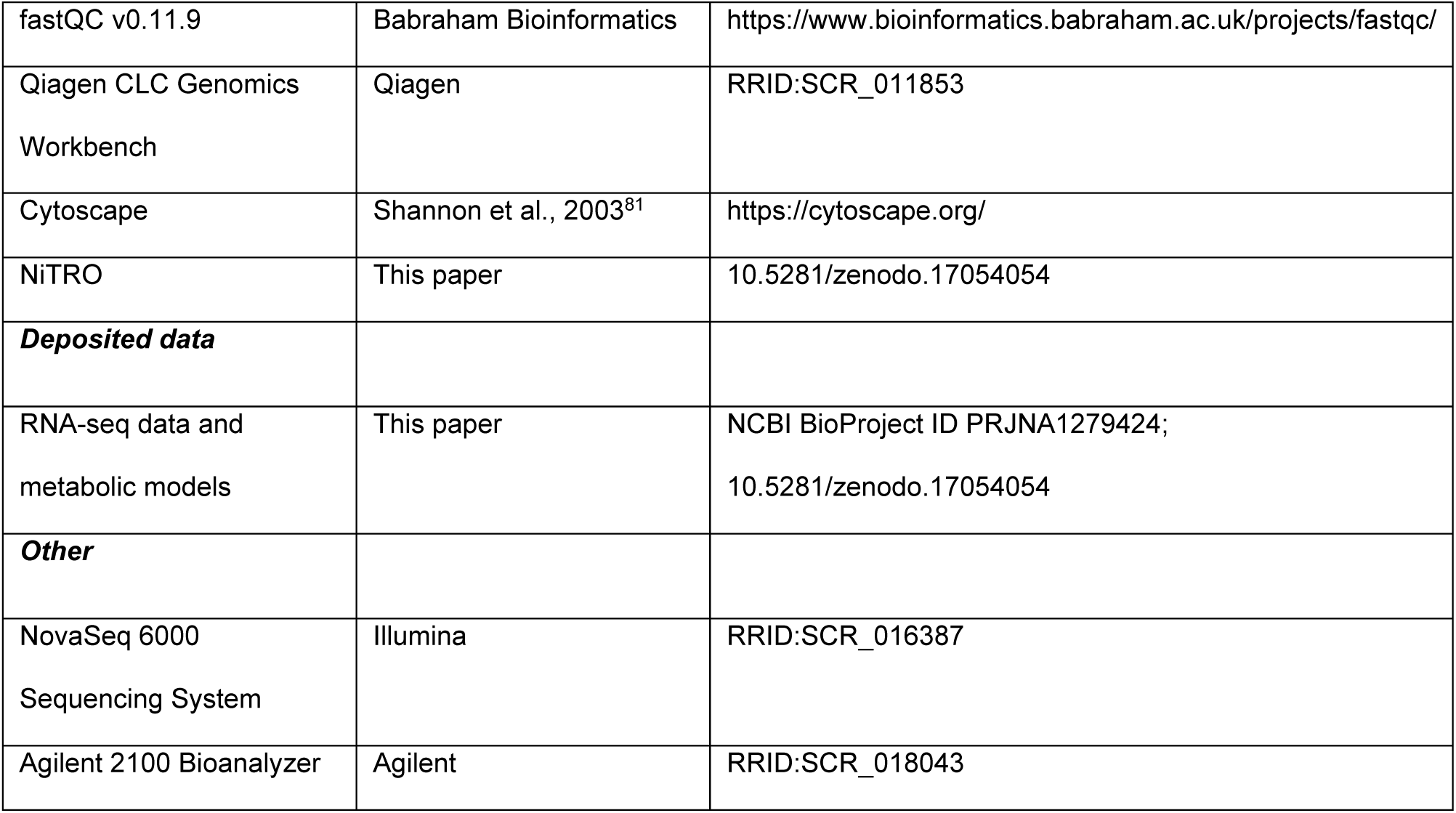

## Supplemental information

Document S1. Figures S1–S7 and Supplementary Tables.

## Acknowledgments

We thank the following GIGA Institute (University of Liege, ULiege) core facilities: Viral Vectors (for BSL3 access and safety control), Genomics (for RNA libraries and sequencing), and Bioinformatics (for data quality control and submission). Computational resources at ULiege have been provided by the Consortium des Équipements de Calcul Intensif (CÉCI), funded by the Fonds de la Recherche Scientifique (FRS-FNRS, Belgium) under Grant No. 2.5020.11 and by the Walloon Region. This work was mainly supported by FRS-FNRS Coronavirus PER-40003579 to J-C.T, and the NYUAD Faculty Research Funds AD060 from the Division of Science (NYUAD, UAE) to K.S-A. J-C. T. is a Research Director of the FRS-FNRS.

Use of AI-assisted tools: ChatGPT (OpenAI) and Perplexity.AI were used to assist in researching published clinical and experimental evidence summarized in Table 1. Claude (Anthropic, Claude Sonnet 4.x, Opus 4.x) was used to assist with manuscript formatting, text editing (including reframing of text, expansion of figure legends, typographical corrections, and figure renumbering), and consistency verification across manuscript and figure files. All AI-generated or AI-assisted content was reviewed, verified, and approved by the authors, who take full responsibility for the accuracy and integrity of the final manuscript.

## Author contributions

B.D. analyzed the data, generated the models and figures and co-wrote the manuscript. D.C.El A. analyzed the data. M.K. analyzed the data. C.P. co-supervised RNA-sequencing. P.F-B. co-supervised the study. J-C.T. Designed wet lab experiments, co-wrote the manuscript and co-supervised the study. D.R.N. co-wrote the manuscript. K.S-A. conceived the study, co-supervised the study, and co-wrote the manuscript.

## Supplemental figure legends

**Figure S1. Volcano plots of DEG for SARS-CoV, CoV-2 and MERS at 24H and 48H**

Volcano plots showing differentially expressed genes (DEGs) in HEK293-ACE2 cells following infection with SARS-CoV (A), SARS-CoV-2 (B), and MERS-CoV (C) at 24 and 48 hours post-infection. Each point represents a single gene; the x-axis shows the log2 fold change in expression relative to mock-infected controls, and the y-axis shows the –log10 adjusted p-value (FDR-corrected). Upregulated and downregulated genes meeting the significance threshold are shown in color; non-significant genes are shown in grey. Related to Figure 1.

**Figure S2. Perturbation on the reaction directionality.**

A reversible reaction is a reaction that has forward and reverse directions. A positive or negative flux value points to the set of metabolites that are produced or consumed, respectively, through the reaction. When the sign of the flux value changes, it indicates alteration on the equilibrium of the reaction. In this figure, we show those reactions (y-axis) that have the same sign in all infected models (boxes), but the opposite sign in the control model (black dots). The x-axis shows the flux value of the corresponding reaction. (A) and (B) correspond to 24H and 48H timepoints, respectively. Related to Figures 4 and 6.

**Figure S3. Significantly enriched SLC family members.**

Enrichment analysis of solute carrier (SLC) family members whose expression is significantly altered following coronavirus infection. Bars represent SLC genes encoding amino acid and metabolite transporters that are enriched among the differentially expressed genes in SARS-CoV-, SARS-CoV-2-, and MERS-CoV-infected HEK293-ACE2 cells at 24 and 48 hours post-infection (FDR < 0.05). Of the 53 annotated SLC25 family members, 21 showed significant expression changes following infection, with distinct temporal dynamics and directionality across the three viruses. Related to Figures 5 and 6.

**Figure S4. The NiTRO algorithm.**

(A) Context-specific genome-scale metabolic models are constructed by integrating RNA-seq expression data from control and coronavirus-infected HEK293-ACE2 cells with the human metabolic reconstruction Recon3D using the GIMME algorithm. (B) Sick reactions are identified by comparing flux ranges between infected and control models using the Jaccard similarity index; reactions with less than 50% flux-range overlap are classified as perturbed. (C) Rescuer gene-pair knockouts are predicted through systematic combinatorial gene-pair deletion in the infected model, followed by partial least squares dimensional reduction to select the top 1% of deletions whose flux profiles correlate linearly with control-model profiles. (D) Reverted reactions are identified by assessing whether post-knockout flux ranges recover more than 50% of the corresponding control-model ranges. Rectangles, circles, and diamonds represent reaction subsets, metabolic analyses, and statistical tests, respectively. Related to Figures 7–9.

**Figure S5. Non-perturbed reactions following infections with coronaviruses**

We use the term “healthy/nonperturbed reaction” when the flux range of the reaction in the infected model covers more than 50% of that in the control model. The top Venn diagrams show the number of nonperturbed reactions at 24H (A) and 48H (B) that are common or exclusive among the infected models. The bottom Venn diagrams show the number of nonperturbed reaction-associated metabolites that are common or exclusive among the infected models. In all panels, red, blue, and light blue shaded circles correspond to SARS-CoV, -CoV-2, and MERS-CoV, respectively. Related to Figure 8.

**Figure S6. Druggable rescuer gene pairs.**

Cytoscape network representation of druggable gene pairs in which both genes are targets for drugs. After assigning the drugs that inhibit the rescuer knockouts, the druggable gene pairs are connected. A rectangle corresponds to a gene that is a partner of a rescuer druggable gene pair. Red and blue edges, correspond to druggable rescuer gene pairs that are predicted in SARS-CoV and SARS-CoV-2, respectively. Related to Figure 9.

**Figure S7. Repurposed drugs.**

A network of interactions between drugs and their target genes, as well as interactions among genes identified as rescuer knockouts. Pink diamonds represent drugs, while light blue circles denote genes corresponding to rescuer knockouts (KOs). The pink edges link each drug to its corresponding target genes, indicating known drug-gene interactions. Light blue edges connect genes that correspond to rescuer KOs, illustrating genetic interactions that play a role in rescuing cells from viral infection. This comprehensive network highlights both drug-target relationships and the gene pairs involved in rescue processes, offering insights into potential therapeutic targets and the genetic landscape of cellular rescue mechanisms. Related to Table 1.

